# A Quantitative Assessment of Prefrontal Cortex in Humans Relative to Nonhuman Primates

**DOI:** 10.1101/233346

**Authors:** Chad J Donahue, Matthew F Glasser, Todd M Preuss, James K Rilling, David C Van Essen

**Author notes:** Correspondence: David C Van Essen: 660 S. Euclid Ave; Box 8108; St. Louis, MO. 63110.

## Abstract

Humans have the largest cerebral cortex among primates. A long-standing controversy is whether association cortex, particularly prefrontal cortex (PFC), is disproportionately larger in humans compared to nonhuman primates, as some studies report that human PFC is relatively expanded whereas others report uniform PFC scaling. We address this controversy using MRI-derived cortical surfaces of many individual humans, chimpanzees, and macaques. We present two parcellation-based PFC delineations based on cytoarchitecture and function and show that a previously used morphological surrogate (cortex anterior to the genu of the corpus callosum) substantially underestimates PFC extent, especially in humans. We find that the proportion of cortical gray matter occupied by PFC in humans is up to 86% larger than in macaques and 24% larger than in chimpanzees. The disparity is even greater for PFC white matter volume, which is 140% larger in humans compared to macaques and 71% larger than in chimpanzees.

## Introduction

Cerebral cortex varies dramatically in size and surface area across mammals. In absolute extent, human cerebral cortex is the largest among primates, with a surface area roughly three-fold larger than in chimpanzees and about 10-fold larger than the intensively studied macaque monkey (1-6). Many studies have reported that association cortex (prefrontal, temporal and parietal regions implicated in higher cognition and affect (7-9)) is disproportionately larger in humans relative to nonhuman primates (10-16). However, other studies have reached opposite conclusions, especially in relation to prefrontal cortex (PFC), resulting in an ongoing controversy (17-19).

A central issue is how the size of PFC scales relative to that of other brain domains. Such analyses are often viewed through the lens of allometry, by comparing the size of a given brain region to some other measure, such as overall brain size, across a range of species. Allometric scaling implies a linear relationship when plotting data on a logarithmically-scaled plot, where the slope of the best-fitting line may show positive (slope > 1; hypermetric), isometric (slope = 1), or negative (slope < 1; hypometric) allometry. Furthermore, a large positive deviation of a single species from an allometric relationship would be referred to as ‘exceptional’ (e.g. human brain size is exceptionally large compared to other mammals (20)). Some previous studies have reported a positive allometric relationship across primate species based on the size of PFC and the rest of the brain using structural volumes (18, 21). Passingham & Smaers (12) similarly reported a positive allometric relationship between volumes of PFC and primary visual cortex (area V1). In contrast, Gabi et al. (17) recently reported an isometric relationship using neuronal counts for PFC vs other cortical regions.

Other morphometric analyses inform but do not resolve this debate. Semendeferi and colleagues (19, 22) reported that although human frontal, temporal and parietal lobes are larger than those of apes, the proportion of total cortex belonging to each region is similar across species. By contrast, studies utilizing cortical surface-based interspecies registration (mapping) driven by putative cortical homologues suggest that surface area in these regions is disproportionately larger (20-fold or more in places) in humans compared to macaques, whereas early sensory regions (e.g. primary visual cortex) are expanded as little as two-fold (3-5). Furthermore, cortical myelin maps derived from MRI (6), reveal a greater extent of lightly myelinated cortex in both association and higher-order sensory regions in humans compared to chimpanzees and macaques, whereas species differences appear more modest in heavily myelinated primary sensorimotor regions (6).

These conflicting results and interpretations regarding PFC scaling are largely attributable to methodological differences between studies. One difference is the region with which PFC is being compared. Bush and Allman (21) compared frontal gray matter with remaining neocortical gray matter, while Smaers et al. (23) made comparisons with more conserved cortical regions such as primary visual cortex (area V1). While some studies have focused on volumetric differences in cortical gray matter and/or extent of the underlying white matter (24), others have considered neuronal counts for gray matter and non-neuronal cells for white matter (17). Most striking, however, are differences in delineating what constitutes PFC. A scarcity of comparative architectonic, anatomical or functional studies that could directly identify the location of homologous areas and regions has led some investigators to instead invoke neuroanatomical proxies for PFC. Semendeferi et al. (19) analyzed the entire frontal lobe, while Schoenemann et al. (24) and Gabi et al. (17) more explicitly approximated PFC using a morphological surrogate: cortex anterior to the genu of the corpus callosum. However, the accuracy of such approximations has not to date been critically assessed.

Generally, PFC has referred to frontal lobe association cortex lying anterior to motor and premotor regions. For over a century, neuroanatomists have attempted to objectively delineate PFC based on cytoarchitectonics. In the primate frontal lobe, cortical layer 4 can appear ‘granular’ (high density of small neurons), ‘agranular’ (lacking a well-defined layer 4) or an intermediate ‘dysgranular’ (subtle layer 4 with a modest density of small neurons). The earliest delineation of PFC was Brodmann’s (15) *regio frontalis*, consisting of granular frontal and orbital cortex. More recent architectonic studies consider PFC to include both granular and dysgranular regions of medial frontal and orbitofrontal cortex in human (25) and macaque (26) and in lateral frontal cortex of both species (27-29). Geyer (30) investigated the functional boundary separating premotor and prefrontal cognitive regions in humans, reporting that agranular cortex in posterior Brodmann area 6 (premotor) transitions anteriorly to dysgranular cortex by a graded cytoarchitectonic transition rather than a sharp boundary. Consistent with this hypothesis, functional neuroimaging analyses implicate some agranular as well as dysgranular frontal lobe areas in higher cognitive function (31).

To address differences in PFC across human and nonhuman primate species, we used structural MRI datasets from humans, chimpanzees, and macaque monkeys to generate cortical surface models of individual subjects, estimate cortical myelin content and thickness, and register individuals to species-average atlases. We then utilized architectonic, functional and neurophysiological data across species to delineate PFC. We present a set of species-specific PFC delineations based on architectonic criteria in humans, chimpanzees and macaques; functional criteria for humans and macaques; and a critical evaluation of the callosal genu PFC approximation (17, 24).

## Results

### Cytoarchitectural designations allow surface-based PFC delineation across species

Based on cytoarchitectonics, we delineated a conservative PFC boundary (Fig. 1, regions filled in red) that includes granular and dysgranular frontal lobe cortex anterior to motor and premotor cortex, as has been proposed for both humans (25, 28) and macaques (32). However, agranular regions of medial frontal and orbitofrontal cortex have been implicated in higher cognitive function, and we consider these appropriate to include in a liberal PFC delineation (Fig. 1, additional regions filled in blue): frontal lobe cortex that is functionally neither motor nor pre-motor.

**Figure 1.**
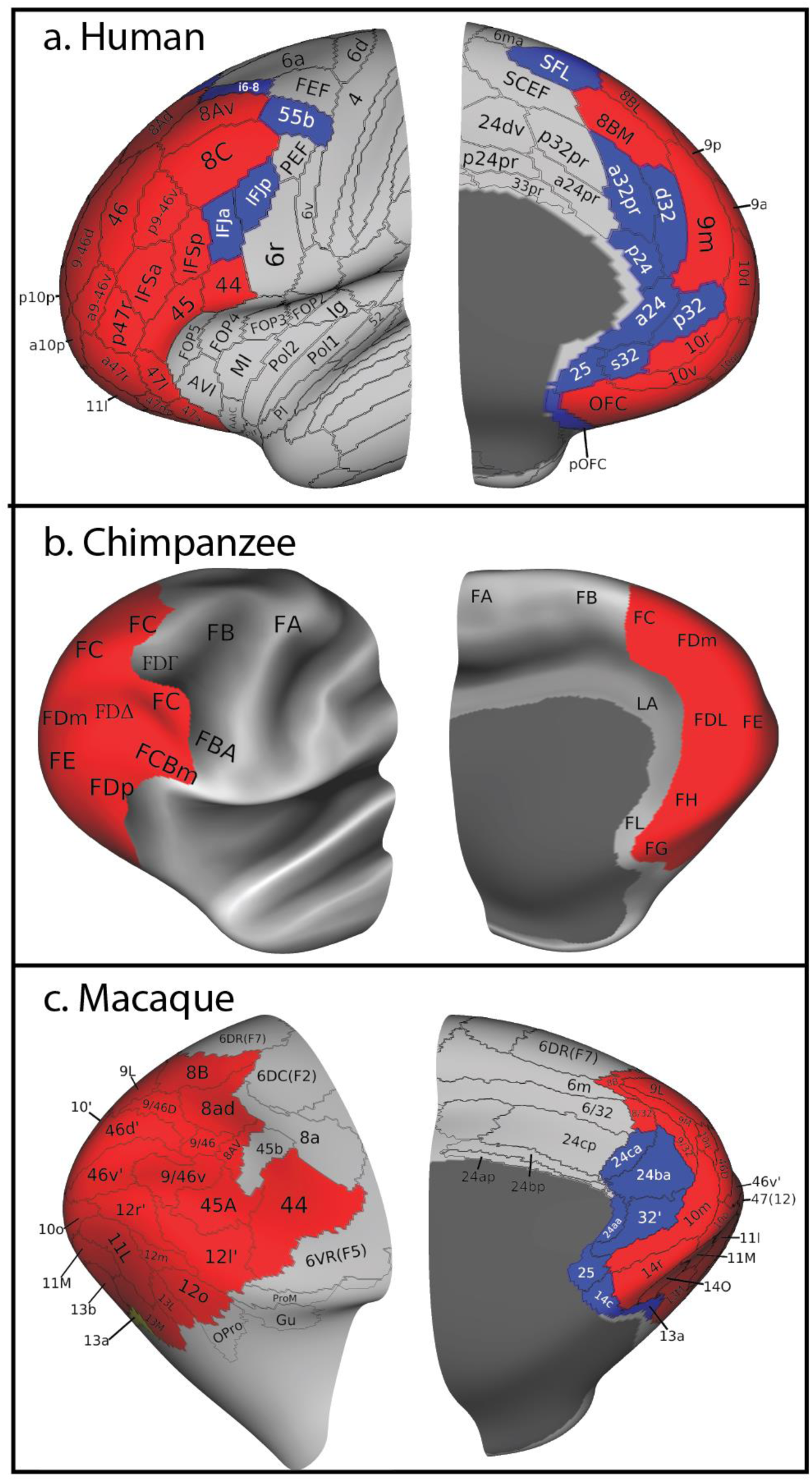
Parcellations of prefrontal cortex for human, macaque, and chimpanzee, displayed on inflated left hemisphere surfaces, cropped to include only anterior regions. A) Inflated (unfolded) human cortical surface displaying group-average HCP_MMP1.0 parcellation. Conservative PFC includes red areas. Liberal PFC additionally includes blue areas. B) Inflated chimpanzee cortical surface displaying a PFC delineation based on the Bailey et al. (33) cytoarchitectonic parcellation plus maps of myelin content. C) Inflated macaque cortical surface displaying a composite parcellation adapted from three studies (34-36) along with conservative and liberal PFC delineations. Figures are not to scale. Data will be made available at https://balsa.wustl.edu/GrK7 (human), https://balsa.wustl.edu/px4G (chimpanzee) and https://balsa.wustl.edu/k94P (macaque) upon acceptance.

For human cortex, we used individual-subject parcellations from 60 unrelated subjects taken from the 210V (‘validation’) group of the HCP Multimodal Parcellation (HCP_MMP1.0), which had been parcellated using an areal classifier that matched individual-subject ‘feature vectors’ to an initial group-average multimodal parcellation (31). For the macaque, a composite of previously published parcellations (34-36) covering the entirety of neocortex was mapped onto a species-average macaque atlas (37) (Yerkes19) and subsequently mapped to the constituent 19 individual subjects for morphometric analyses. This composite includes several new or modified areas, reflecting a combination of previously described areas (marked as ‘prime,’ e.g., area 10’ found in dorsolateral PFC is an amalgamation of area 10 as reported by Ferry et al. (34) and Paxinos et al. (36)), with some undergoing further subdivision. Specifically, areas 24a, 24b and 24c were subdivided into anterior and posterior segments (e.g. 24aa and 24ap, respectively), based on descriptions of area 24 as dysgranular anteriorly with associated cognitive-related function transitioning to an agranular cytoarchitecture posteriorly associated with motor function (27, 38, 39). The chimpanzee architectonic parcellation of Bailey et al. (33) includes areal designations on a series of histological section contours. We mapped the areal designations from these section contours to corresponding MRI sections in an individual chimpanzee brain and from the individual to our surface-based chimpanzee atlas (see SI). We then drew estimated areal boundaries on the atlas surface, aided in dorsolateral cortex by myelin gradients described in the next section.

For each species, an area was considered part of conservative PFC based on cytoarchitecture and inclusion in previously published PFC delineations. Areas unique to the human liberal PFC delineation include IFJa/p, SFL, i6-8 and 55b of lateral frontal cortex, a/p24, 25, d/p/s32 and a32pr of medial frontal cortex and posterior orbitofrontal cortex (pOFC). For the macaque, exclusively liberal PFC areas include medial frontal areas 24aa, 24ba, 24ca, 25 and 32’, and orbitofrontal areas 13a and 14c. The frontal eye fields (areas FEF and PEF in humans, area 45b in the macaque) are sometimes considered part of PFC (38, 40), but we excluded them here because of their stronger association with premotor regions than with cognitive regions in terms of moderate rather than sparse myelin content (see below) and their functional connectivity (31). Because modern functional data is unavailable for the chimpanzee, we focused only on a conservative PFC delineation based on presence of granular/dysgranular frontal lobe cytoarchitecture. We delineated Bailey et al. agranular areas as motor (FA), premotor (FB and FBA), and cingulate (LA and FL). More anterior regions were designated as PFC, except that we excluded FDr from PFC based on its being moderately myelinated (see below) and its likely involvement in eye movement control. Supporting Tables S1 & S2 provide additional information for human and macaque candidate PFC areas, including references for areal cytoarchitecture and cognitive-related designations.

### Structural cortical features aid in delineating PFC

Maps of estimated myelin content (based on the T1w/T2w intensity ratio (6, 41)) provide a useful architectural marker for identifying cortical regions and areas across species, particularly in the chimpanzee where cytoarchitectural data are limited and functional data is unavailable. Spatial gradients of these myelin maps can provide objective evidence of sharply defined architectonic transitions (e.g. from dense to moderate or light myelination- see (41)). Figure 2 illustrates myelin content and the corresponding spatial gradient for each species in relation to our three PFC delineations. These reveal important patterns and correlations across measures, but the relationship to PFC boundaries is complex and differs for dorsolateral, ventrolateral, and medial regions.

**Figure 2.**
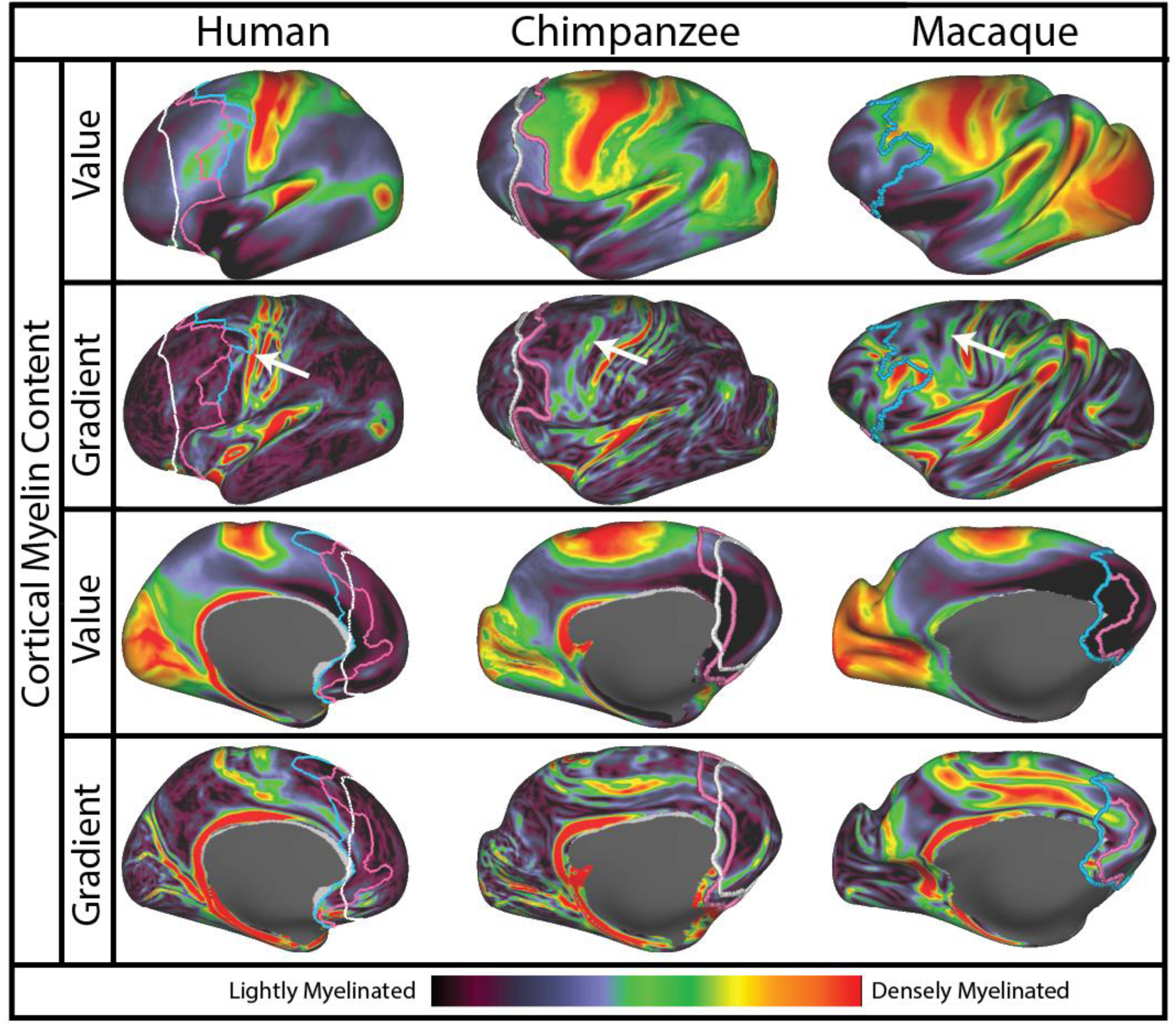
Structural cortical features related to PFC delineations. Left hemisphere frontal cortex of each primate species displaying myelin content (first and third rows) and its corresponding spatial gradient (second and fourth rows). The white line overlying each map represents the group-average location of the coronal slice at the corpus callosum genu (see also Fig. 3); pink and blue lines represent group-average conservative and liberal PFC delineations, respectively. Data will be made available at https://balsa.wustl.edu/22XL upon acceptance.

In dorsolateral cortex, the heavily myelinated primary somatomotor strip (top row, in red) and the moderately myelinated premotor strip (green in human and chimpanzee, mainly orange-yellow in macaque) provide useful landmarks across species. In all three species, a myelin gradient ridge (white arrows, second row) runs along the transition between premotor and primary motor strips. A second, more subtle, gradient ridge lies anteriorly. In the human, this anterior gradient ridge aligns well with the liberal PFC border. In the macaque, both conservative and liberal PFC borders run in the general vicinity of the anterior myelin gradient ridges, but the correlation is not good for either. This gradient ridge informed our delineation of chimpanzee conservative PFC, whose border was drawn to follow the ridge closely in this region.

In ventrolateral and ventral regions, lightly myelinated PFC in all three species is adjoined by even more lightly myelinated cortex in the anterior insula (top row, indigo and black) and orbitofrontal cortex (third row). However, the most prominent myelin gradient does not coincide with published architectonic PFC delineations (also, in the macaque and chimpanzee datasets, orbitofrontal cortex was not as accurately segmented owing to localized signal dropout and distortion of T1w relative to T2w images). In medial cortex, PFC is lightly myelinated dorsally and very lightly myelinated ventrally in all three species, but none of them show a clear myelin gradient running along the PFC boundary. Thus, there are strong cross-species similarities in myelin maps in and near PFC, but only in dorsolateral PFC of human and chimpanzee do we consider the myelin gradients to be strongly informative about PFC borders. We also compared PFC boundaries to cortical thickness maps and their gradients. As with the myelin maps, there are some correlations, but they are of limited utility for delineating PFC borders (see SI Fig. 1).

### Genu-based morphological surrogate underestimates PFC extent

Figure 3 illustrates the range of individual variation in the location of liberal, conservative, and genu-based PFC borders as defined for each species. Probability maps indicate most (yellow) and least (black) common locations of each border

**Figure 3.**
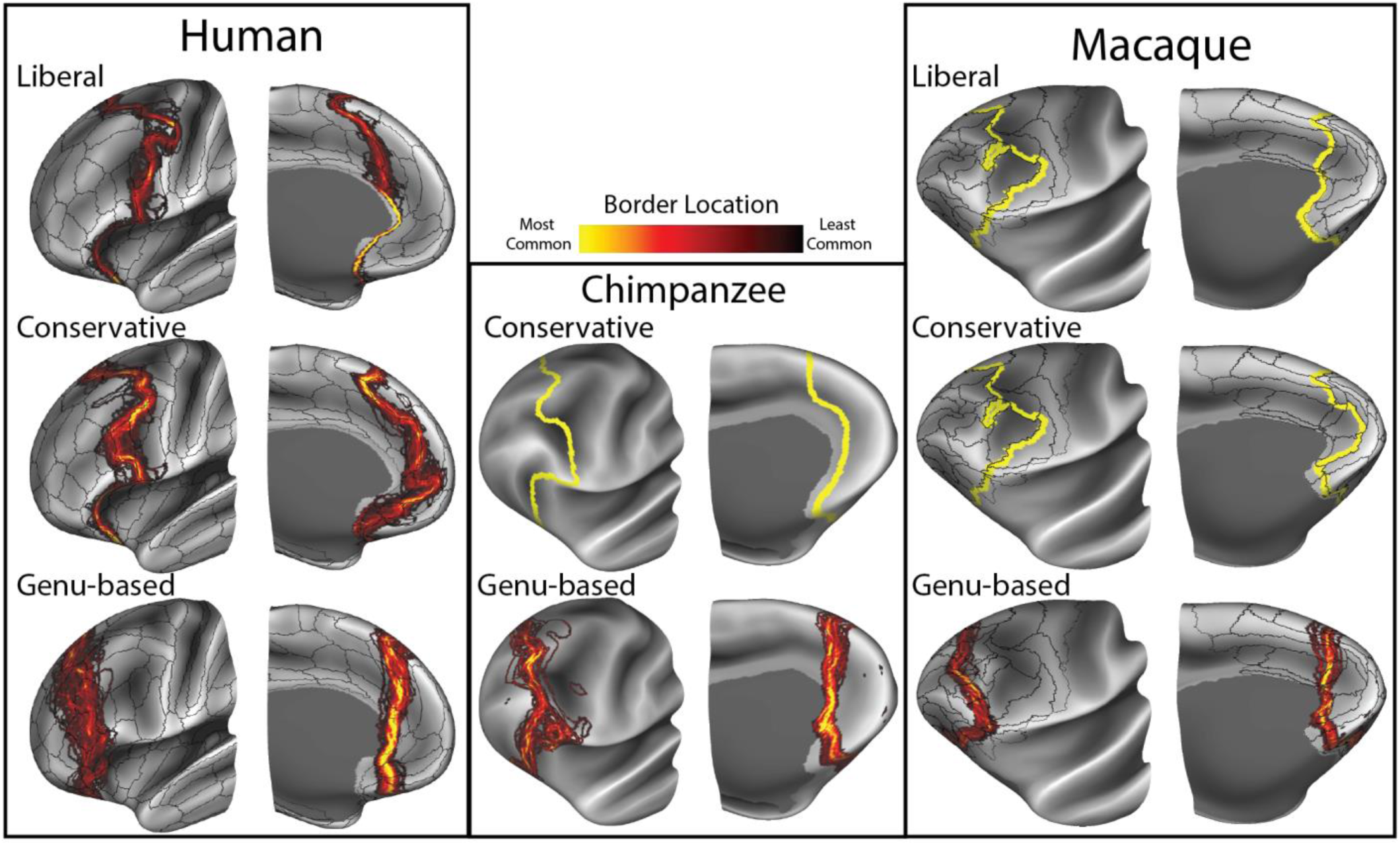
PFC border probability maps displayed on inflated atlas surfaces of the frontal lobe. Human liberal and conservative PFC borders were created using individual subject parcellations, resulting in intersubject variance that becomes evident when displayed on the group-average surface. Corresponding borders in the macaque and chimpanzee were created using a group-average parcellation registered to each individual subject, and thus shows no such variance. Probability maps were created based on PFC border in individual subjects (human n=60; macaque n=19; chimpanzee n=29) and are displayed on group-average surfaces with underlying group-average sulcal depth maps. Data will be made available at https://balsa.wustl.edu/r7Xw (human), https://balsa.wustl.edu/xMp4 (chimpanzee) and https://balsa.wustl.edu/PMKk (macaque) upon acceptance.

Important species differences are revealed by comparing parcellation-based PFC delineations (both liberal and conservative) to genu-based PFC approximations (Fig. 3, third row). In dorsolateral frontal cortex, the macaque and chimpanzee genu-based PFC border runs somewhat anterior to the moderately myelinated region adjacent to premotor cortex, mostly anterior to the conservative delineation. By visual inspection, the genu-based delineation in the human more substantially underestimates PFC spatial extent when compared to both liberal and conservative delineations. This observation is evaluated quantitatively in the next section.

### Human PFC is absolutely and relatively large compared to nonhuman primates

Table 1 reports the mean volume for total cortical gray matter, primary visual cortex area V1 and mean volumes relating to PFC delineations, along with standard deviations. These volumes were calculated for both hemispheres of each individual subject and then averaged. Also provided (in parentheses) is the percentage of each PFC ROI volume and of area V1 relative to total volume.

**Table 1.** Volumes of human, chimpanzee and macaque cortical gray matter for entire cortex; genu-based, conservative and liberal delineations of association cortex; and primary visual area V1. Percentages in parentheses are of total volume.

| **Gray Matter**<br>**(All values cm<sup>3</sup>)** | **Total Volume** | **Genu-based PFC** | **Conservative PFC** | **Liberal PFC** | **Area V1** |
| --- | --- | --- | --- | --- | --- |
| **Human**<br>(n = 60) | 512 ± 55 | 77.2 ± 11 (15%) | 105 ± 13 (21%) | 131 ± 16 (26%) | 14.0 ± 2.0 (3%) |
| **Chimpanzee**<br>(n = 29) | 134 ± 13 | 18.5 ± 2.6 (14%) | 23.2 ± 2.4 (17%) |  | 7.5 ± 0.8 (6%) |
| **Macaque**<br>(n = 19) | 34.4 ± 2.9 | 3.34 ± 0.5 (10%) | 4.4 ± 0.5 (13%) | 4.8 ± 0.6 (14%) | 3.7 ± 0.4 (11%) |

After averaging the left and right hemispheres, computed mean gray matter volumes were 513 cm^3^, 134 cm^3^ and 34.4 cm^3^ for human, chimpanzee and macaque, respectively, indicating that human cortical gray matter is 15-fold greater value than the macaque and roughly 4-fold greater than the chimp. Mean surface areas were 1,843 ± 196 cm^2^ for humans, 599 ± 53 cm^2^ for chimpanzees and 193 ± 13 cm^2^ for macaques; mean cortical thicknesses were 2.69 ± 0.09 mm, 2.63 ± 0.09 mm and 2.02 ± 0.08 mm, respectively (standard deviations are of means across subjects). Thus, the species differences in volume ratios (15-fold vs macaque; 4-fold vs chimpanzee) exceeds that for the corresponding surface areas (10-fold vs macaque; 3-fold vs chimpanzee) and thickness (1.3-fold vs macaque; 1.02-fold vs chimpanzee) measurements.

As shown by our two parcellation-based delineations, the proportion of PFC gray matter volume is up to 1.9- fold larger in humans compared to macaques (26% vs 14% for the liberal delineation; 21% vs 13% for the conservative delineation) and 1.2-fold larger compared to chimpanzees (21% vs 17% for the conservative delineation). The genu-based PFC approximation shows a more moderate species difference, with PFC**genu** constituting 15% of human cortical gray matter compared to 14% in the chimpanzee and 10% in the macaque. Cortical volume computed using the genu-based proxy for PFC underestimates the parcellation-based delineations in all three species, but less so in macaques and chimpanzees and more so in humans.

#### Scaling relative to area V1

To quantify how PFC has scaled relative to a more evolutionarily conserved cortical region, we analyzed primary visual cortex (area V1) in terms of its volume, surface area and cortical thickness in humans and macaques. To identify area V1, we used the HCP_MMP1.0 (31) delineation of V1 in the human, the Lewis & Van Essen (35) delineation in the macaque, and myelin maps plus the Bailey et al. (33) delineation in the chimpanzee (see SI). In humans, chimpanzees and macaques, respectively, we found mean V1 volume was 14.0 cm^3^, 7.5 cm^3^ and 3.7 cm^3^; surface area was 69.7 cm^2^, 47.2 cm^2^ and 24.1 cm^2^; and cortical thickness was 1.99 mm, 2.01 mm and 1.76 mm. While V1 is comparable in size to PFC in the macaque, the volume of PFC gray matter is up to 3-fold larger than area V1 in the chimpanzee and up to 9-fold larger in humans. Surface area followed a similar trend (PFC up to 1.8-fold more than V1 in chimpanzees and 6-fold in humans). Cortical thickness of PFC and V1 scaled similarly across species.

#### White matter volumes

Table 2 reports for each species the total volume of subcortical white matter along with volumes of white matter anterior to the genu-based PFC proxy, which is the only parcellation having a well-defined posterior extent of PFC white matter and hence amenable to analysis of white matter volumes. Total white matter volumes for humans, chimpanzees and macaques were on average 443 cm^3^, 119 cm^3^ and 21.8 cm^3^, respectively.

**Table 2.**
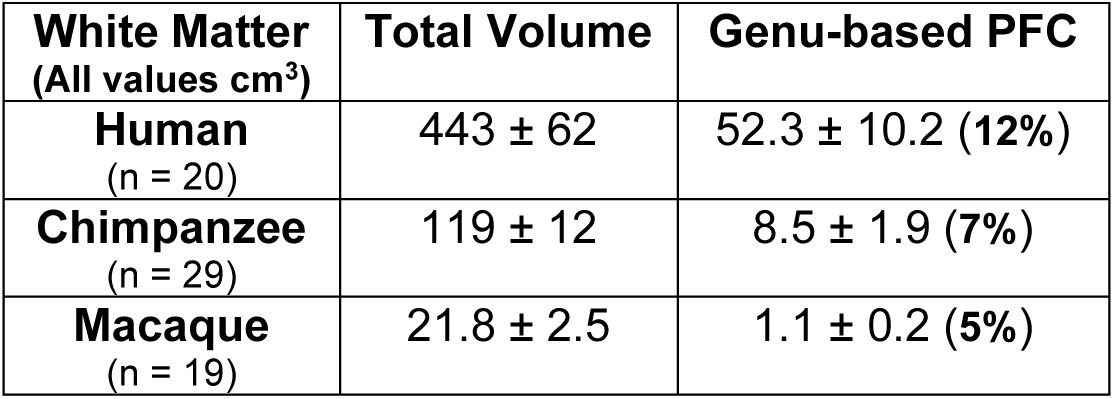
Volumes of human, chimpanzee and macaque subcortical white matter for entire cortex and genu-based delineations of association cortex (see Methods). PFC percentages in parentheses are relative to total volume.

| White Matter<br>(All values cm<sup>3</sup>) | Total Volume | Genu-based PFC |
| --- | --- | --- |
| **Human**<br>(n = 20) | 443 ± 62 | 52.3 ± 10.2 (12%) |
| **Chimpanzee**<br>(n = 29) | 119 ± 12 | 8.5 ± 1.9 (7%) |
| **Macaque**<br>(n = 19) | 21.8 ± 2.5 | 1.1 ± 0.2 (5%) |

The ratio of total white matter to gray matter volume is similar in the human and chimpanzee (0.86 & 0.89, respectively), considerably greater than that for the macaque (0.63), indicating a 35 – 40% relative increase of total white matter to gray matter ratio in humans and chimpanzees relative to the macaque. Genu-based PFC white matter to gray matter ratios are 0.68 in the human, 0.46 in the chimpanzee and 0.34 in the macaque. Analyzed differently, PFC white matter is a larger fraction of total white matter in humans (12%) than in chimpanzees (7%) or macaques (5%). This 2.4-fold relative difference in PFC white matter between macaques and humans (12% vs 5%) markedly exceeds the 1.5-fold difference in the genu-based gray matter volumes.

### PFC exhibits positive allometric scaling for humans and nonhuman primates

Figure 4 illustrates several logarithmically-scaled comparisons that describe the relative size increase of PFC across species to reference measures. In order to assess the differential scaling of different cortical regions, we plotted volumes of PFC gray matter, non-PFC gray matter and primary visual cortex against the volume of total cortical gray matter (Fig. 4A). We additionally compared PFC gray matter to the more evolutionarily conserved primary visual cortex (Fig. 4B) and PFC white matter to total white matter (Fig. 4C).

**Figure 4.**
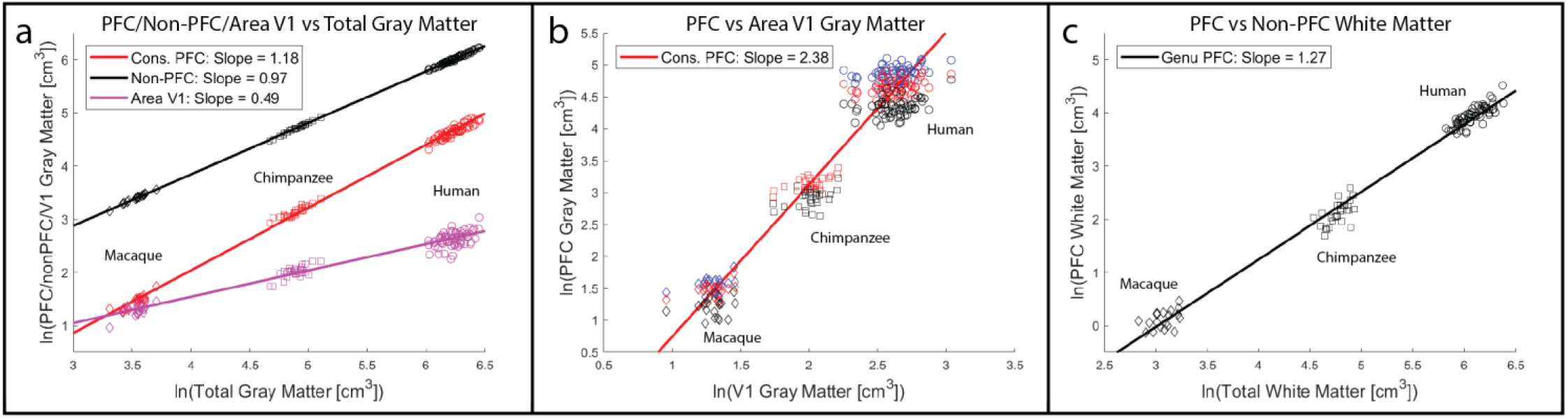
Log-scale plots comparing PFC gray and white matter volume to reference values for genu-based (black), conservative (red) and liberal (blue) PFC delineations. A) Volumes of conservative PFC, non-PFC and area V1 gray matter plotted against total cortical gray matter volume. B) PFC gray matter volume plotted against primary visual cortex (area V1) gray matter volume. Blue, red and black markers indicate liberal, conservative and genu-based PFC delineations, respectively. C) Genu-based PFC white matter volume plotted against total white matter volume. For all panels, solid lines represent best fit using mean macaque, chimpanzee and human data points.

When using total cortical gray matter as a reference (Fig. 4A), the scaling of conservative PFC gray matter as defined by the regression line through the mean macaque, chimpanzee and human data points exhibits positive allometry (slope of 1.18) which is notably steeper than the regression lines for non-PFC cortex (slope of 0.97) and for area V1 (slope of 0.49). When instead using the evolutionarily-conserved visual area V1 as a reference (Fig. 4B), PFC scales with even greater positive allometry (slope of 2.38). Finally, PFC white matter scaled with total white matter volume with positive allometry (slope of 1.27).

## Discussion

Using a cortical surface-based approach, we have presented a comparative delineation and analysis of frontal association cortex in humans, chimpanzees and macaques. In addressing a longstanding controversy, we report strong evidence for a greater proportion of human PFC gray matter volume compared to two nonhuman primates and an even greater species difference for PFC white matter volume.

### Prefrontal cortex can be delineated using cytoarchitectonic and functional criteria

Our criteria for areal inclusion in PFC entailed judgment calls based on the preponderance of available evidence. Our conservative criteria relating to granular/dysgranular cytoarchitecture, although guided by evidence in the literature, is not a simple consensus view. Our liberal delineations in the human and macaque (areas reported to have agranular cytoarchitecture) entailed subjective assessments regarding what cognitive task contrast activation or other functional information relative to neighbors warrant inclusion. A notable exception to these criteria across species is the exclusion of frontal eye fields: area FEF in humans, 45b in macaques and FDr in chimpanzees. While these areas are cytoarchitecturally granular/dysgranular and receive thalamic inputs primarily from the mediodorsal (MD) nucleus (see below), their functional relationship to attentional eye movement led to our placing them in the ‘motor’ category as opposed to cognitive-related PFC. Additionally, some areas in the macaque composite parcellation (e.g. anterior and posterior subdivisions of areas 24a, 24b and 24c) might reasonably be reassigned to account for a gradient in cytoarchitecture, primarily in areas that extend posteriorly into densely myelinated cortex. However, plausible alternative choices for PFC extent in any of the species would not negate our main conclusion that the relative size of PFC in the human lineage is larger than that in nonhuman primates. The lack of functional imaging data in chimpanzees limited our analysis to conservative and genu-based PFC delineations largely informed by the Bailey et al. cytoarchitectural atlas (33) and maps of myelin content. Our candidate chimpanzee PFC boundary between premotor and prefrontal cortex warrants further analysis using modern architectonic and/or imaging methods.

Besides the architectonic, functional, and morphological criteria invoked in our study, other types of information have been proposed for delineating PFC in various mammalian species. These include the distribution of dopaminergic projections and a prominent level of projections from the mediodorsal (MD) nucleus of the thalamus (42-44). Both metrics, however, have not been shown to be adequately specific to provide strong diagnostic criteria for delineating PFC (29, 38): dopaminergic projections are prominent throughout the primate brain and are not demonstrably overrepresented in granular frontal cortex (45, 46) compared to more posterior neocortex. Similarly, MD thalamic projections, while particularly prevalent in granular frontal cortex and thus a meaningful guide, are also present across the precentral motor region as well as more posterior regions (47).

### Human PFC volume is disproportionately large

We used a surface-based approach (1, 2, 4) derived from structural MRI to generate our analysis of cortical volumes. Our sample size includes 60 humans, 29 chimpanzees and 19 macaques, thereby enabling estimates of variability in each population. We found an increase in the proportion of PFC cortical gray matter volume in humans up to 1.9-fold compared to macaques and up to 1.2-fold compared to chimpanzees. The increase in underlying white matter volume is even more pronounced with volume in humans up to 2.4-fold larger than macaques and 1.7-fold larger than chimpanzees. Thus, consistent with several previous studies (2, 5, 6, 24), our results strongly support the hypothesis that the proportion of cortical gray and white matter volumes attributed to frontal association cortex is differentially larger in humans compared to nonhuman primates. Furthermore, the absolute size of conservatively delineated human PFC is 4.7-fold larger than in the chimpanzee, which is especially striking considering the more modest difference in size of primary visual cortex (1.9-fold larger in humans). Coupled with the evidence that humans have substantially more PFC white matter volume, this points to an impressively greater amount of neural machinery associated with PFC in humans compared to nonhuman primates.

Though other studies have argued for a more general scaling-up of the human brain compared to nonhuman primates (17, 19), we do not view the idea of a predictive positive allometric scaling as mutually exclusive from preferential expansion of PFC or association cortex in general. For example, Semendeferi et al. (19) reported that the proportion of the cortical mantle occupied by cortex anterior to the precentral gyrus does not differ significantly between humans and great apes. However, Passingham and Smaers (12), analyzing the size of various cortical regions relative to primary visual cortex (used as a reference), found a positive allometric scaling of association cortex (their Figure 1 & Table 1). Our analyses corroborate those from Passingham and Smaers by revealing a positive allometric relationship of PFC gray matter volume when comparing to total gray matter volume (Fig. 4A; best-fit line slope of 1.18) and even more so when compared to primary visual cortex (Fig. 4B; slope of 2.38). However, we note that the individual slope values reported for the regressions in this study are limited by the inclusion of only three species. Therefore, we emphasize how the regression slopes differ when comparing different regions of cortex: when using total cortical gray matter as a reference, conservative PFC exhibits positive allometry compared to non-PFC and particularly area V1. Furthermore, the non-PFC delineation still contains highly expanded regions of association cortex in the parietal and temporal lobes, thus biasing its scaling toward positive allometry (compared to the negative allometric scaling reported for area V1 gray matter vs total cortical gray matter).

Defining PFC as anterior to the genu of corpus callosum, Gabi et al. (17) used an isotropic fractionator approach (48) to count neurons and non-neuronal cells as well as gray matter and white matter volumes as measured in 2 mm coronal tissue slabs through one hemisphere in a single brain from each of 8 primate species. For humans vs the average of two macaque species, they reported 10% vs 7.6% of cortical gray matter, 5.5% vs 4.5% of white matter, and 8% vs 7.35% of total cortical neurons belonging to genu-based PFC (their Table S1). Thus, for these two species, their results suggest a slightly positive allometric relationship for PFC cortical gray matter and a nearly isometric relationship for neuronal numbers. In the present study, we found a greater species disparity for both genu-based (15% vs 10%) and parcellation-based (21% vs 13%) delineations. Our finding that the genu-based approximation underestimates PFC volume (to somewhat different degrees in humans, chimpanzees and macaques) (Fig. 4) is consistent with the prediction of Schoenemann et al. (24).

Along with their allometric analyses, Gabi et al. reported that human neuronal density increases anteriorly to posteriorly with lowest density at the frontal pole, whereas macaques exhibit a ‘double gradient’ of high neuronal densities near both the frontal and occipital poles. A complementary perspective comes from cellular neuroanatomical analyses by Elston and colleagues (49-52), revealing that pyramidal neurons in nonhuman primate granular PFC (and in other regions of association cortex such as lateral temporal cortex) exhibit more complex dendritic structure (size of the dendritic tree, branching structure and spine density) when compared to the less elaborate dendritic structure found in evolutionarily conserved regions such as sensorimotor and early visual cortex (51). Thus, the differential expansion in PFC volume documented in the present study likely in part reflects increased synaptic ‘machinery’ and not simply an increased number of PFC neurons in the human lineage.

### White matter underlying human PFC is particularly large compared to nonhuman primates

We found that human PFC white matter volume (as a percentage of total white matter) is 2.4-fold greater than in the macaque and nearly two-fold greater than in the chimpanzee (Table 2, Fig. 4C). It is intriguing to speculate on the neuroanatomical basis of these striking species differences. Given the aforementioned evidence (17) that PFC neuronal density is relatively low in humans vs NHPs, the density of output axons from human PFC projection neurons (i.e., pyramidal cells) is presumably lower as well. A disproportionately large white matter volume underlying human PFC might instead reflect (i) an increased density of afferent projections from distant (non-PFC) regions and contributing to an increased axonal density within human PFC gray matter; (ii) a disproportionately high percentage of human PFC output axons that traverse the underlying white matter but nonetheless terminate within other PFC targets; (iii) a disproportionately large average axonal diameter in human PFC white matter; and/or (iv) a disproportionately high degree of axonal branching within human PFC white matter. Disentangling these and other possibilities is unlikely to be easy but might become feasible with further advances in postmortem neuroanatomical methods.

### Improving the granularity of interspecies comparisons

Our analysis has focused on measurements of PFC in its entirety, even though it is very heterogeneous in its internal organization, connectivity, and function. Previous comparisons between macaque and human that used interspecies surface-based registration (4, 5) provided evidence that relative expansion in the human lineage is highly nonuniform within PFC as well as in other higher cognitive regions (e.g. lateral parietal and temporal cortex) relative to early sensory and motor regions. However, such “evolutionary expansion” maps should be interpreted with caution, given that (i) some of the candidate homologues are plausible but not firmly established and (ii) the surface-based registration algorithm used to constrain the interspecies mapping tolerated local nonuniformities that are not well-grounded neurobiologically. Recent algorithmic improvements in surface-based registration such as the Multimodal Surface Matching method (53, 54) should help address the latter problem when adapted to interspecies registration constraints. The former problem (identifying candidate homologues) should benefit from recent advances in parcellating human cortex (31) and characterizing its network organization (especially resting-state networks), combined with recent and prospective advances in parcellating macaque cortex and characterizing its network organization.

## Methods

### Data Collection

Healthy, adult human structural T1-weighted (T1w) and T2-weighted (T2w) scans were acquired at 0.7 mm isotropic resolution as part of the Human Connectome Project (HCP), using the HCP’s standard protocol (55). From a larger HCP subject set, 60 unrelated subjects (30 male and 30 female) were selected for analysis from the S500 HCP data release. Macaque and chimpanzee structural T1w and T2w scans were acquired at the Yerkes National Primate Research Center at Emory University. A group of 19 adult macaques (1 male and 18 female) were scanned at 0.5 mm isotropic resolution. A group of 29 adult chimpanzees (all female) were scanned at 0.8 mm isotropic resolution. For the macaque and chimpanzee datasets, localized signal dropout was observed in anterior insular and orbitofrontal cortex.

### Image Preprocessing

For each human subject, T1w and T2w scans were initially processed using the minimal preprocessing pipelines developed for the HCP (55) to maximize alignment across imaging modalities and minimize distortions & blurring of the data. Field maps were available and readout distortion was corrected in humans. Intersubject registration to a group-average atlas surface was performed using a two-stage process based on the multimodal surface matching (MSM) algorithm (53), where an initial gentle stage is driven by cortical folding patterns and a more aggressive second stage utilizes cortical areal features of myelin, resting-state network maps, and visuotopic maps (31).

A version of the HCP pipelines adapted for nonhuman primates (HCP-NHP pipelines (37)) was used to process macaque and chimpanzee T1w and T2w structural scans. Initially, the PreFreeSurfer pipeline aligns T1w and T2w volumes to native anterior commissure-posterior commissure (AC-PC) space and performs brain extraction, cross-modal registration, bias field correction and nonlinear volume registration to atlas space using the FMRIB Software Library (FSL (56)).

A nonhuman primate-specific pipeline, FreeSurferNHP differs from the HCP FreeSurfer pipeline in the following ways: (1) using nonhuman primate volume templates and adjusting the brain size parameter to 80 mm for the macaque and 120 mm for the chimpanzee and (2) converting the data into a “fake” 1 mm isotropic 256^3^ space to conform to FreeSurfer requirements without interpolation (57). This last modification allows the full resolution of the non-human primate data to be used for surface generation. Species-specific volume and surface templates were used for registration prior to transformation back into the 0.5 mm input space. Cortical thickness is then computed, modifying the maximum cortical thickness parameter (from a default value of 5 mm) in the FreeSurfer *mris_make_surfaces* command to conform to scan resolution as 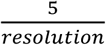 mm to ensure that the fake 1 mm space does not set too low a cap on the cortical thickness (0.5 mm data faked to 1 mm would make a 5 mm cortical thickness limit into a 2.5 mm cortical thickness limit if not adjusted).

The PostFreeSurfer pipeline uses MSM-Sulc surface registration, a version of MSM where alignment is driven by cortical shape (FreeSurfer’s “sulc” measure). The pipeline produces a high-resolution ‘164k’ surface mesh (~164,000 vertices per hemisphere), as well as two lower-resolution meshes (32k/10k for macaque and 32k/20k for chimpanzee). In addition to surface reconstruction, myelin maps (average maps of myelin content across the cortical layers) are created based on the T1w/T2w ratio (6, 41, 55). However, the cortical thickness and myelin maps are likely biased in regions of localized signal dropout (anterior insular and orbitofrontal cortex), because of an imperfect cortical segmentation.

### Group-Average Atlases & Cortical Parcellations

Individual subject registration to a group-average atlas enables accurate comparisons across subjects. The HCP Multimodal Parcellation (HCP_MMP1.0 (31)) provided both an atlas space for registration of individual human subjects and a cortical areal classifier for creating individual subject parcellations. This classifier identified the 180 cortical areas per hemisphere if the fingerprint of each area was detected, but allowed for identification of fewer areas. These areas vary in size and shape relative to the original areas defined using group average data. Additionally, a more accurate medial wall was created to restrict analyses to neocortex and transitional cortex (but excluding the hippocampal formation medial to the presubiculum).

The Yerkes19 macaque surface-based atlas (37) was created using the 19 previously described adult macaque subjects. The HCP-NHP pipelines were used to extract cortical surfaces and subcortical volumes from structural MRI scans. Interhemispheric alignment was driven by 45 geographically corresponding landmark contours per hemisphere, analogous to the landmark-based alignment performed for the F99 macaque surface-based atlas (2). The Ferry et al. (34), Lewis & Van Essen (35) and Paxinos et al. (36) histological parcellations were mapped to the Yerkes19 atlas from the F99 atlas (34, 36, 58) using the MSM algorithm driven by a combination of cortical folding (mean curvature) and revised medial walls (excluding hippocampal cortex medial to the presubiculum) for both atlases. Mean curvature was used as a registration constraint because an accurate FreeSurfer-based ‘sulc’ map was not available for the F99 atlas surface, which had been generated using the SureFit segmentation method (3) that does not produce the white matter surface required to create a ‘sulc’ map.

The Yerkes29 chimpanzee surface-based atlas was created in a similar fashion, using the 29 previously described adult chimpanzee subjects. The Bailey et al. (33) cytoarchitectural atlas was used to systematically map cortical areas from published coronal volume slices to the Yerkes29 atlas. Images of histological slice drawings (intermediate between coronal and axial planes) taken from the Bailey et al. atlas were visually matched to corresponding slices in an individual chimpanzee MRI structural scan, using an individual (Edwina) whose frontal convolutions were similar to the atlas, based on visual inspection. These coronal areal designations were then projected to the cortical group average surface based on Bailey et al.’s cortical surface figures and our group-average maps of myelin and sulcal depth.

Delineations approximating PFC based on the genu of the corpus callosum were created by identifying the coronal slice precisely anterior to the genu when individuals were aligned so that the axial slice was parallel to the AC-PC line. All gray and white matter anterior to this coronal slice was considered part of this genu-defined region.

### Assigning Prefrontal Cortical Areas

Individual cortical areas were identified as belonging to PFC based on published criteria and delineations (25-27, 33). For each cortical area located in the frontal lobe, primary qualities tabulated included previously published areal classification as PFC and the histological description of areal cortical layer IV (granular, dysgranular, lightly granular or agranular). Cytoarchitectonically granular/dysgranular areas were included in the delineation of conservative PFC, while agranular areas associated with cognitive-related function were additionally included in the liberal PFC delineation. Supporting text and Tables S1-S2 provide additional information about the studies used to define PFC delineations.

### Calculation of Cortical Gray Matter and White Matter Volumes

Total cortical volumes were determined by isolating the cortical gray matter ribbon (as defined by the space between white and pial surfaces) and the underlying white matter (as defined by FreeSurfer segmentation (57, 59)) in the native subject space. Total cortical surface areas and mean cortical thicknesses were computed for each subject using the native midthickness surface mesh and excluding the medial wall, using a revised medial wall demarcation for the macaque relative to a published version (37) (by exclusion of hippocampal cortex and other minor adjustments). For each *conservative* and *liberal* parcellation-based PFC delineation, constituent areas were adjoined to create contiguous PFC surface-based regions of interest (ROI). In humans, these ROIs were created on each subject’s 32k mesh using each subject’s individual HCP_MMP1.0 parcellation and subsequently registered to each subject’s native space. In macaques, these ROIs were created on a 164k mesh using the composite parcellation defined on the “Yerkes19” group-average surface, and subsequently registered to each subject’s native surface mesh. Similarly, the chimpanzee ROI was created on the “Yerkes29” 164k surface mesh and mapped to native subject meshes. Genu-based ROIs were created by including all surface vertices rostral to a coronal slice of the AC-PC aligned volume at the genu of the corpus callosum, an approximation of PFC used in previous studies (17, 24). Volume measurements were determined by summing the volumes of individual polyhedral “wedges” within each ROI, where each wedge is defined by a triangle in the “white” surface and the corresponding triangle in the pial surface. This process was performed on each individual subject (human n=60; macaque n=19; chimpanzee n=29) and the mean and standard deviation of all subjects were reported for each case (Table 1). This process is illustrated for and exemplar human in Figure 5.

**Figure 5.**
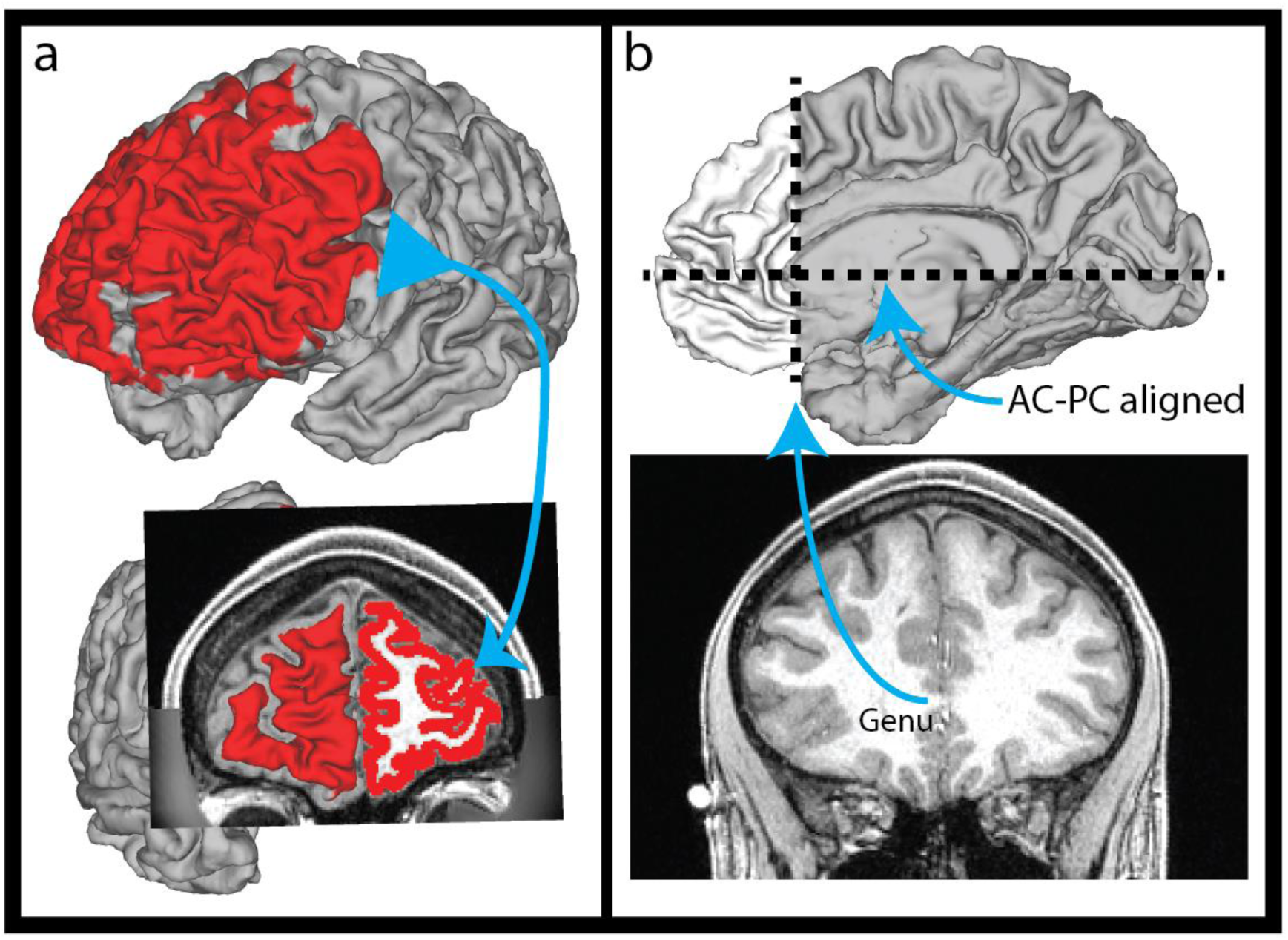
Mapping of surface-based ROI to cortical gray matter ribbon volume. A. Illustration of human cortical surface ROI mapped to underlying gray matter ribbon volume. B. Illustration of human genu-based ROI volume anterior to coronal slice at genu of the corpus callosum when image is AC-PC aligned.

## Data Availability

All data related to this study as well as supporting information will be freely available upon acceptance via the BALSA database (https://balsa.wustl.edu/) at https://balsa.wustl.edu/study/zlVX.

## Acknowledgments

We thank T. Coalson and E. Reid for their aid in completing this study. This work was supported by grants NIH 5T32EB01485506 (CJD), NIH F30 MH097312 (MFG) and RO1 MH-60974 (DCVE). Human datasets were provided by the Human Connectome Project, WU-Minn Consortium (Principal Investigators: David Van Essen and Kamil Ugurbil; 1U54MH091657) funded by the 16 NIH Institutes and Centers that support the NIH Blueprint for Neuroscience Research; and by the McDonnell Center for Systems Neuroscience at Washington University. Macaque and chimpanzee datasets were provided through support from National Institutes of Health Grant P01AG026423 and National Center for Research Resources P51RR165 (superseded by the Office of Research Infrastructure Programs/OD P51OD11132), and by the National Chimpanzee Brain Resource, R24NS092988.

